# Ketamine selectively enhances AMPA neurotransmission onto a subgroup of identified serotoninergic neurons of the rat dorsal raphe

**DOI:** 10.1101/2020.08.19.255794

**Authors:** Cyrine Hmaied, Stanislav Koulchitsky, Ivan Gladwyn-Ng, Vincent Seutin

## Abstract

Although the fast antidepressant effect of ketamine is now well established clinically, neither its mechanism(s) nor its main site(s) of action is clearly defined. Because enhanced serotoninergic (5-HT) transmission is an important part of the antidepressant effect of various drug classes, we asked whether ketamine and one of its metabolites (hydroxynorketamine [HNK]), both used in their racemic form, may modulate the excitatory drive onto these neurons.

Using whole-cell recordings from pharmacologically identified 5-HT and non-5-HT neurons in juvenile rat dorsal raphe slices, we found that both ketamine and HNK (10 µM) increase excitatory AMPA neurotransmission onto a subset (50%) of 5-HT neurons, whereas other 5-HT cells were unaffected. Both compounds increased the amplitude as well as the frequency of spontaneous excitatory post-synaptic currents (sEPSCs) mediated by AMPA receptors. The effect of ketamine was more robust than the one of HNK, since it significantly enhanced the charge transfer through AMPA channels, whereas HNK did not. The increase in the excitatory drive induced by ketamine was dependent on NMDA receptor blockade. In the presence of tetrodotoxin, the effect of ketamine was markedly reduced. Non-5-HT neurons, on the other hand, were unaffected by the drugs.

We conclude that ketamine and HNK increase the excitatory drive onto a subset of 5-HT neurons by promoting glutamate release and possibly also through a postsynaptic action. The effect of ketamine is dependent on NMDA receptor modulation and appears to involve a network effect. These findings improve our understanding of the fast-acting antidepressant effect of ketamine.

**SIGNIFICANCE STATEMENT:** The mechanisms of ketamine’s antidepressant effect are currently controversial. We asked whether the drug would produce changes is the strength of synaptic inputs onto serotoninergic neurons of the dorsal raphe. We found that this is indeed the case in about half of these neurons. The action of ketamine was mimicked to some extent by its well-known metabolite hydroxynorketamine, was dependent on NMDA receptor activation and probably involved a local network effect. It remains to be determined if the differential susceptibility of serotoninergic neurons to the drug correlates with any differential inputs and/or outputs.

## INTRODUCTION

Several randomized placebo-controlled clinical studies have now established the usefulness of a low-dose intravenous (i.v.) injection of the non-competitive NMDA antagonist ketamine as a fast-acting antidepressant therapy (Newport et al., 2015), although the exact indications for its use are still a matter of debate (Sanacora et al., 2017). Very recently, intranasal administration of its S-enantiomer was shown to be effective in depressed patients who are resistant to conventional antidepressants (Daly et al., 2018) and was approved by the FDA as a therapeutic option in this population.

Since 15 years, numerous studies have attempted to circumscribe the mechanisms of action of ketamine, with a number of controversial results. Globally, the following facts are more or less established: the acute antidepressant effect of ketamine may not entirely depend on its NMDA antagonism because other NMDA antagonists do not share this action (Zanos et al, 2016, but see Yang et al., 2018). One or several metabolites (in particular 2R-6R hydroxynorketamine [HNK]) probably contribute to this effect, perhaps by acting on a different target (Zanos et al., 2016, but see Suzuki et al., 2017). Evidence suggests that the drug potentiates excitatory AMPA-mediated glutamate neurotransmission and increases synaptic strength in various brain areas. The mechanisms by which ketamine produces this effect are debated. Several possibilities exist (reviewed in Zanos and Gould 2018). The « disinhibition hypothesis » posits that the drug preferentially inhibits inhibitory interneurons in the cortex, thereby indirectly increasing the excitability of pyramidal neurons (as shown for MK-801 by Homayoun and Moghaddam, 2007). Others have shown that spontaneous glutamate release from pyramidal neurons is able to activate postsynaptic NMDA receptors at rest. This tonically activates eukaryotic elongation factor 2 (eEF2) via its phosphorylation by its kinase, leading to inhibition of BDNF translation and mTORC1-mediated protein synthesis. By blocking these NMDA receptors, ketamine may upregulate this pathway (Autry et al., 2011) and increase synapse formation (Li et al., 2010). Yet another hypothesis is that extrasynaptic, tonically activated NMDA receptors are responsible for the inhibition of the mTORC1 pathway and that the block of these receptors by ketamine is involved in its therapeutic action.

One other major potential target for ketamine could be the forebrain projecting serotonergic (5-HT) neurons of the dorsal raphe (DR) (du Jardin et al., 2016), since extensive pharmacological, preclinical, clinical and genetic evidence suggests a prominent role for serotonin hypofunction in the genesis of depressive disorders. So far, the hypothesis that ketamine enhances excitatory transmission onto 5-HT neurons has not been tested thoroughly (see Discussion). Using patch clamp recordings in acute rat brain slices, we found that the drug strongly enhances AMPA neurotransmission onto a subgroup of pharmacologically identified DR 5-HT neurons, whereas it does not affect inputs onto non-5-HT (presumably GABAergic) neurons in this region.

## MATERIALS AND METHODS

### Ethical approval

Experiments were performed in accordance with the national Belgian laws concerning animal experimentation (December 23, 1998 and September 13, 2004), and the European Union (Directive 2010/63/EU). In addition, all animal care, handling and sacrifice was approved by the Ethics Committee for Animal Use of Liège University (protocol 2022).

### Drugs

(R, S)-ketamine hydrochloride was obtained from Pr. Monbaliu (Center for Integrated Technology and Organic Synthesis, Department of Chemistry, Liège University, Belgium). Hydroxynorketamine hydrochloride (HNK) was obtained from Sigma-Aldrich (St. Louis, MO, USA). 8-hydroxy-2-(di-n-propylamino) tetralin, (8-OH-DPAT), 6-cyano-7-nitroquinoxaline-2, 3-dione (CNQX) and (2S)-3-[[(1S)-1-(3, 4-dichlorophenyl) ethyl]amino-2-hydroxypropy](phenylmethyl) phosphinic acid hydrochloride (CGP 55845) were obtained from Biotechne (Minneapolis, MN, USA). D-(-)-2-Amino-5-phosphonopentanoic acid (D-AP5) and tetrodotoxin citrate were purchased from Hello Bio. Gabazine (SR95531) was purchased from Alomone (Jerusalem, Israel). All drugs were stored at −20°C as stock solutions in Milli-Q water, except CGP55845 and CNQX, which were initially dissolved in dimethyl sulfoxide (DMSO) as stock solutions, stored at −20°C and then diluted 1000× in the artificial cerebro-spinal fluid (ACSF, see its composition below) on the day of the experiment. Repetitive control experiments have shown that 0.1% DMSO does not affect 5-HT neuron excitability. Other drugs were diluted to the appropriate final concentration in ACSF as well.

### Brain slices

Experiments were performed on slices from juvenile (3-4 week old) male and female Wistar rats. All animals had been group housed and maintained on a constant 12‐h light–12‐h dark cycle. Brain slice preparation was done between 10 and 11 AM (between ZT 3 and 4) in most cases. The rats were anesthetized using chloral hydrate (400 mg⁄kg, i.p.). After decapitation, brains were rapidly removed and placed in ice-cold (0–4°C) ACSF bubbled with carbogen (95% O_2_, 5% CO_2_). The ACSF was composed of (in mM): 125 NaCl, 2.5 KCl, 1 MgCl_2_, 2 CaCl_2_, 1.25 NaH_2_PO_4_, 10 glucose, 25 NaHCO_3_, had an osmolarity of ~310 mOsm/l, and a pH of 7.4. Coronal slices comprising the dorsal raphe nucleus (DRN) (thickness: 300 µm) were cut using a vibroslicer (DTK‐1000; Dosaka, Tokyo) and subsequently incubated in oxygenated ACSF at 34 °C for ~25 min. The slices were next incubated in ASCF at room temperature for at least 30 min before being transferred to the recording chamber where they were completely submerged in the ACSF (flow rate ~2.0 ml/min, temperature 34 °C). When switching solutions, the time for complete exchange of the bath solution was about 1 min. Only one neuron was recorded per slice. In most cases, recordings were obtained in 2-3 slices per animal.

### Whole-cell patch-clamp recordings

Patch pipettes were pulled from thick-walled borosilicate glass capillaries (2.0 mm outer diameter, 0.5 mm wall thickness; Hilgenberg, Malsfeld, Germany) using a Sutter P-87 puller (Sutter Instruments, Novato, CA, USA) and had resistances between 3–6 MΩ when filled with the internal solution (in mM: K-gluconate, 35; KCl, 110; HEPES, 10; MgCl_2_, 2; Na_2_-ATP, 2; GTP-Na_2_, 0.3; EGTA, 1; Na_2_-phosphocreatine, 10; (osmolarity ~300 mOsm/l, pH adjusted to 7.2 with KOH). A K^+^-based internal solution was chosen in order to be able to check for the presence or absence of the 5-HT_1A_-induced outward GIRK current. We are aware that this solution distorted to a significant extent the postsynaptic currents located at distant synapses. However, this was not a problem in this study because we did not aim for a very precise biophysical characterization of these events. Moreover, the persistence of the dendritic filtering allowed us to evaluate whether or not the ketamine-induced events were located at a similar electrotonic distance from the soma as compared to the spontaneous events. Under our conditions, both spontaneous excitatory and inhibitory post-synaptic currents (sEPSCs and sIPSCs) could be recorded as inward currents at −60 mV.

Signals were amplified and voltage or current commands were applied using a Multiclamp 700B amplifier connected to a Digidata 1440A interface. They were stored using pClamp 10.4 software (Molecular Devices, Sunnyvale, CA, USA). The signals were digitally sampled at 10 kHz and subsequently filtered at 2 kHz for analysis. The reference electrode was an Ag–AgCl pellet immersed in the bath solution. Possible liquid junction potentials were neglected. Only cells with a stable resting membrane potential ≥ −50 mV and with an input resistance higher than 150 MΩ were accepted for further measurements. The access resistance was < 15 MΩ in all cells and experiments in which the access resistance varied by more than 20% during the experiment were discarded.

Neurons were visualized with an Olympus BX51W microscope fitted with a 40X water-immersion objective and equipped with an ICD-42B Ikegami camera. In current clamp, putative 5-HT neurons had electrophysiological characteristics (long duration action potential, large and long-lasting afterhyperpolarization, lack of sag during a hyperpolarizing current pulse) corresponding to those previously described by various groups, including ours (Alix et al., 2014; Calizo et al, 2011). However, it has been shown that several parameters from immunohistochemically identified 5-HT neurons and non 5-HT neurons at least partially overlap (Calizo et al, 2011). Because our recordings were long-lasting, post-hoc immunohistochemical identification of recorded neurons proved unreliable in preliminary experiments. Therefore, we defined the neurochemical nature of recorded neurons by their response to the selective 5-HT_1A_ receptor agonist 8-OHDPAT (100 nM). These receptors are considered to be mostly present in 5-HT neurons in the dorsal raphe. 5-HT neurons were defined as those in which 8-OHDPAT elicited a reversible outward current of at least 30 pA at −60 mV in voltage clamp mode. Using this cut-off value allowed us to exclude some non-5HT neurons which have been reported to undergo a small hyperpolarization with the 5HT_1A_ agonist 5-CT (Beck et al., 2004). The mean outward current for the whole population of 5-HT neurons was 72.0 ± 3.7 pA (n = 39). In most experiments, recordings were obtained from the ventromedial part of the DRN (vmDRN), where the 5-HT neurons are the densest (Ren et al., 2018).

### Protocols

Glutamate receptor–mediated spontaneous excitatory post-synaptic currents (sEPSCs) were recorded in voltage clamp. To isolate them, recordings were performed in the continuous presence of SR 95531 (10 µM) and CGP 55845 (1 µM). A stable baseline was obtained for 10 minutes during the application of the blockers. Ketamine (10 µM) or (2R, 6R)-HNK (10 µM) were subsequently bath applied over a period of 30 minutes. After that period and a 10-minute wash-out period, the slice was superfused with 8-OHDPAT to determine the nature of the recorded neuron. The glutamatergic nature of the sEPSCs was verified at the end of the experiment by applying the glutamate receptor antagonists D-AP5 (50 μM) and CNQX (10 µM). In some experiments, the AMPA nature of the recorded sEPSCs was confirmed by their suppression by CNQX alone. In one set of experiments; we tested whether NMDA receptor blockade by ketamine was responsible for its effects on AMPA sEPSCs. To do so, we added 50 µM APV to SR95531 and CGP 55845 throughout the experiment. The same strategy was applied when we examined the influence of TTX.

### Data analysis

The Mini Analysis software (Synaptosoft, version 6.0.7) was used to analyze synaptic currents. In order to characterize control synaptic transmission, we generally used for analysis the last 3 min before ketamine/HNK superfusion. In most experiments, we compared sEPSC parameters of this period and those recorded during the last 3 minutes of the ketamine/HNK superfusion period. In order to evaluate the time course of the effect of ketamine and HNK, we also measured sEPSCs parameters during the last minute of the control period and during every other minute of superfusion of ketamine/HNK (see Fig. 3C).

The sEPSC events were selected individually and separated from the artifacts using the area and amplitude thresholds. sEPSCs with multiple peaks, which were rather rare, were excluded from the analysis.

Two sets of data were generated. In the first set, we used a threshold of −10 pA and visually selected all events that had the shape of a typical AMPA EPSC. This set was used to analyze the frequency of the events. In the second set, the detection threshold was set at −20 pA for the analysis of the amplitude as well as the total charge transfer (Q). In this case, all the detected events were aligned, superimposed and averaged to obtain one single average trace. Q was calculated for a given period as the AUC (in pA.ms) of the averaged EPSC, as given by the software, multiplied by the sEPSC frequency, as measured in this set of data. From this second set of data, we selected events in which the fitting proposed by the software was perfectly accurate (in terms of time of onset, time of peak and shape of decay). This subset was used to determine the rise time (10-90% of peak) and time constant of decay (90-37%) of the events, assuming a single exponential.

### Statistical analysis

Data are presented as mean ± SEM (normal distributions) or median values (non-normal distributions), as determined by a Kolmogorov-Smirnov test. Statistical analyses were performed with GraphPad Prism version 8.0.2 (GraphPad Software, La Jolla, CA). Distributions passing the test for normality were compared using Student’s t test or a one-way ANOVA in the case of comparisons between > 2 groups. In the latter case, the F statistic is presented as F (DFn, DFd). The post-hoc test was the Tukey test as recommended by GraphPad Prism°. We did not perform a nested ANOVA analysis for our results because neurons in slices from different animals have very equivalent properties. Distributions that did not pass this test were compared using nonparametric tests, either the Wilcoxon/Mann-Whitney tests or the Friedman test in the case of comparisons between > 2 groups.

The data in figure 3C on the time course of the effect of ketamine is presented as a so-called violin plot, which was preferred to a box plot because it better describes the distribution of the data. In this figure, one outlier value was detected and deleted using the Rout detection method.

Part of the data is presented as cumulative probability plots. In order to evaluate whether there was a difference between the distribution of the various parameters in control conditions and in the presence of ketamine or HNK, we quantified the equivalent of an EC_50_ value for each cell in the two conditions. This value was the value of the current amplitude (in the case of current amplitude distributions) or the timing (in the case of interevent interval plots) of the event which had 50% of the maximal value. A chi^2^ test was also used in appropriate cases. The level of significance was set at a *p* value of 0.05.

## RESULTS

### sEPSCs are observed in 5-HT neurons of the DRN

In the presence of GABA receptor blockers (10 µM SR95531 and 1 µM CGP 55845 for GABA_A_ and GABA_B_ receptors, respectively), voltage clamp recordings at –60 mV showed the presence of spontaneous fast synaptic events which occurred at a frequency of 2.25 ± 0.32 Hz (n = 30 cells) and had an amplitude of −10 to −50 pA in pharmacologically identified (see Methods) 5-HT neurons (Figs. 1,4,5,6 and 8). These sEPSCs were clearly due to AMPA receptor activation because they were fast (decay time constants (90-37 %) were 2.0 ± 0.09 ms) and completely blocked by 10 µM CNQX (Fig. 1C).

**Figure 1.**
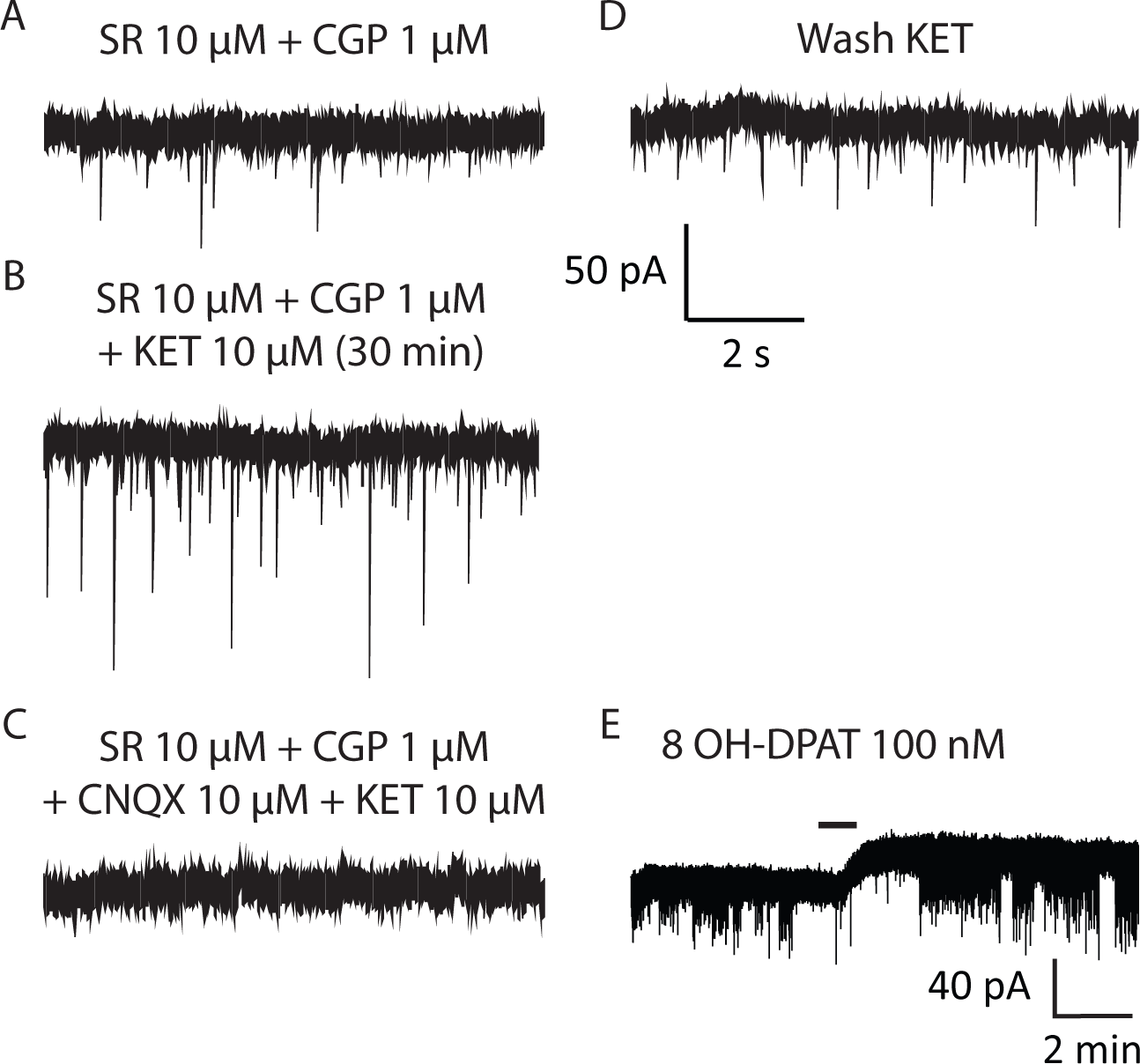
Ketamine increases the amplitude and frequency of AMPA excitatory synaptic currents in a subset of pharmacologically identified 5-HT neurons of the DRN. A, B, Representative traces of sEPSCs recorded in the absence (A) and in the presence (B) of ketamine 10 µM in a 5-HT neuron at a holding potential of −60 mV. C, the sEPSCs is blocked by CNQX (10 µM). D, Representative trace of sEPSCs after washout of ketamine and CNQX. (E) The outward current elicited by the 5HT_1A_ agonist 8-OH-DPAT (100 nM) demonstrates the serotoninergic nature of the recorded neuron.

### Effects of ketamine on sEPSCs in 5-HT and non-5-HT neurons

We examined the effect of 10 µM ketamine on sEPSCs in 5-HT and non-5-HT neurons of the ventromedial portion of the dorsal raphe (DR). Representative traces of sEPSCs from individual neurons are shown for 5-HT neurons in Figs. 1 and 4.

Application of 10 µM ketamine produced a progressively developing and robust increase of AMPA sEPSCs in some, but not all, pharmacologically identified 5-HT neurons within the dorsal raphe (Figs. 1 and 4). We defined neurons as responsive to the drug when both the mean amplitude and the mean frequency of the events were increased by at least 20%. Using these criteria, 8 out of 16 neurons were found to be responsive to the drug. The detected events were aligned, superimposed and averaged to obtain one single average trace. Average traces of sEPSCs recorded from an individual neuron were used to calculate sEPSC amplitude, rise time (10-90 % in ms and decay time constant (90-37 % in ms) using a monoexponential function. Overall the amplitude of the sEPSCs was increased by ~60% (Fig. 2G; from 28.7 ± 1.5 to 46.3 ± 4.3 pA). When comparing the amplitude of the sEPSCs in control, in ketamine and after washout of the drug, we found that there was a significant difference between the three conditions (one-way ANOVA: F _(1.603, 11.22)_ = 11.90, *p* = 0.0025). A Tukey post-hoc test showed that the amplitude was significantly different in control and ketamine (*p* = 0. 0029, n = 8). On the other hand the amplitude of the sEPSCs was not significantly different between the ketamine superfusion period and its wash-out, suggesting a very partial reversibility (*p* = 0.1292). In these neurons, the median frequency of the events was also increased by ketamine from 2.10 to 4.41 Hz. A Friedman test demonstrated a significant difference between the control, ketamine and wash (*p* = 0.0003, n = 8). Dunn’s post hoc tests showed that the frequency was significantly different in control and ketamine (*p* = 0.0014). In addition, the frequency of sEPSCs almost completely recovered to the baseline level during the wash-out period (from 4.41 to 2.16 Hz, *p*= 0.03).

**Figure 2.**
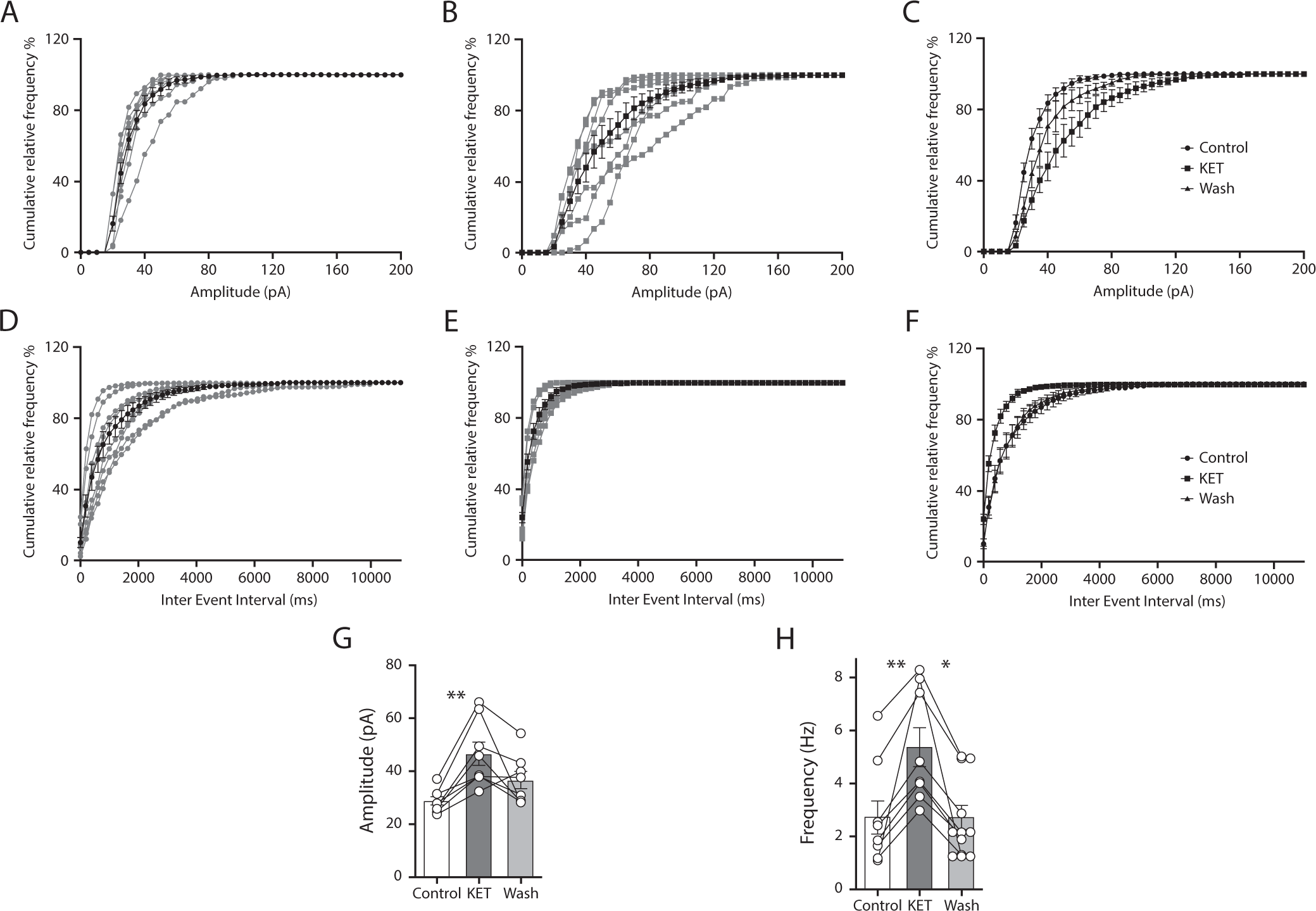
Quantification of the effect of ketamine. A, B, Cumulative probability plots of sEPSC amplitude of the individual cells in control (A), and during ketamine (B) (Bin width = 5 pA, n = 8). C, Cumulative probability plots from all the cells were averaged to generate cumulative probability plots in each condition (control, ketamine, and washout) D, Cumulative probability of sEPSC interevent intervals of the individual cells in control (D) and during ketamine (E) (Bin width = 200 ms).C, Histogram of the averaged data for the amplitude (one-way ANOVA: F (1.603, 11.22) = 11.90, *p* = 0.0025). Tukey post-hoc test showed that the amplitude was significantly different in control and ketamine (*p* = 0. 0029, n = 8). On the other hand the amplitude of the sEPSCs was not significantly different between the ketamine superfusion period and its wash-out (*p* = 0.1292). F, Corresponding average probability plots. G, Histogram of the averaged data for the amplitude. H, average frequency of the sEPSCs

On the other hand, the drug did not affect kinetic parameters of the sEPSCs. The rise time was 0.75 ± 0.02 and 0.89 ± 0.07 ms, in control conditions and in ketamine, respectively (t = 2.09 with df = 7, *p* = 0.074, Student’s t test). The time constant of decay of the events was also unchanged by the drug. Median values were 2.28 and 2.58 ms in control conditions and in ketamine, respectively (Wilcoxon test: *p* = 0.14).

To evaluate the population of sEPSCs more precisely, cumulative probability plots from sEPSCs recorded in individual neurons were constructed and subsequently averaged (Fig. 2A, B, C). In ketamine-responsive 5-HT neurons, this cumulative probability plot of the sEPSCs amplitude displayed a significant displacement to the right in the presence of the drug. The mean “EC_50_” value (see Methods) rose from 25 to 33.75 pA (Fig. 2C). A Friedman test showed that there was a significant difference between these values in the 3 groups (control, ketamine and wash) *p* = 0.0023, n = 8). Dunn post hoc tests showed a significant difference between control and ketamine (*p* = 0.0081), but not between ketamine and wash (*p* = 0.78). Fig. 2G displays a summary histogram of the effect of ketamine on the amplitude of the events. The cumulative probability distribution of the interevent-intervals was left shifted in comparison to the control and the washout periods (Fig. 2D, E, F). The “EC_50_” value decreased from 600 to 100 ms. A Friedman test showed that there was a significant difference between these values in the 3 groups (control, ketamine and wash), *p* = 0.0054, n = 8). Dunn post hoc tests showed a significant difference between control and ketamine (*p* = 0.026) and no significant difference between ketamine and wash (*p* = 0.18). Fig. 2H displays a summary histogram of the effect of ketamine on the frequency of the events.

The time course of the effect of ketamine was variable from cell to cell (Fig. 3C), but always much longer than the one needed to reach the peak effect of 8-OH-DPAT (~2 minutes: e.g. Fig. 1E. A Kruskal Wallis test showed a significant increase in terms of amplitude over time (*p* = 0.0014, n = 8). A Dunn post hoc test showed a significant effect starting at 12 min (*p* = 0.0047). It is apparent from Fig. 3C that it took in general about 14 minutes for ketamine to produce its maximal effect (*p* = 0.0003 at this time point). The effect lasted until 30 min (*p* = 0.0009 at this time point). On the other hand, the drug did not modify the holding current at −60 mV, suggesting that it did not have an effect on the intrinsic excitability of 5-HT neurons (holding current was −13.75 ± 7.30 and −12.5 ± 6.19 pA at baseline and during ketamine (t=1 with df = 7, *p* = 0.3506, paired Student’s t test, n = 8).

**Figure 3.**
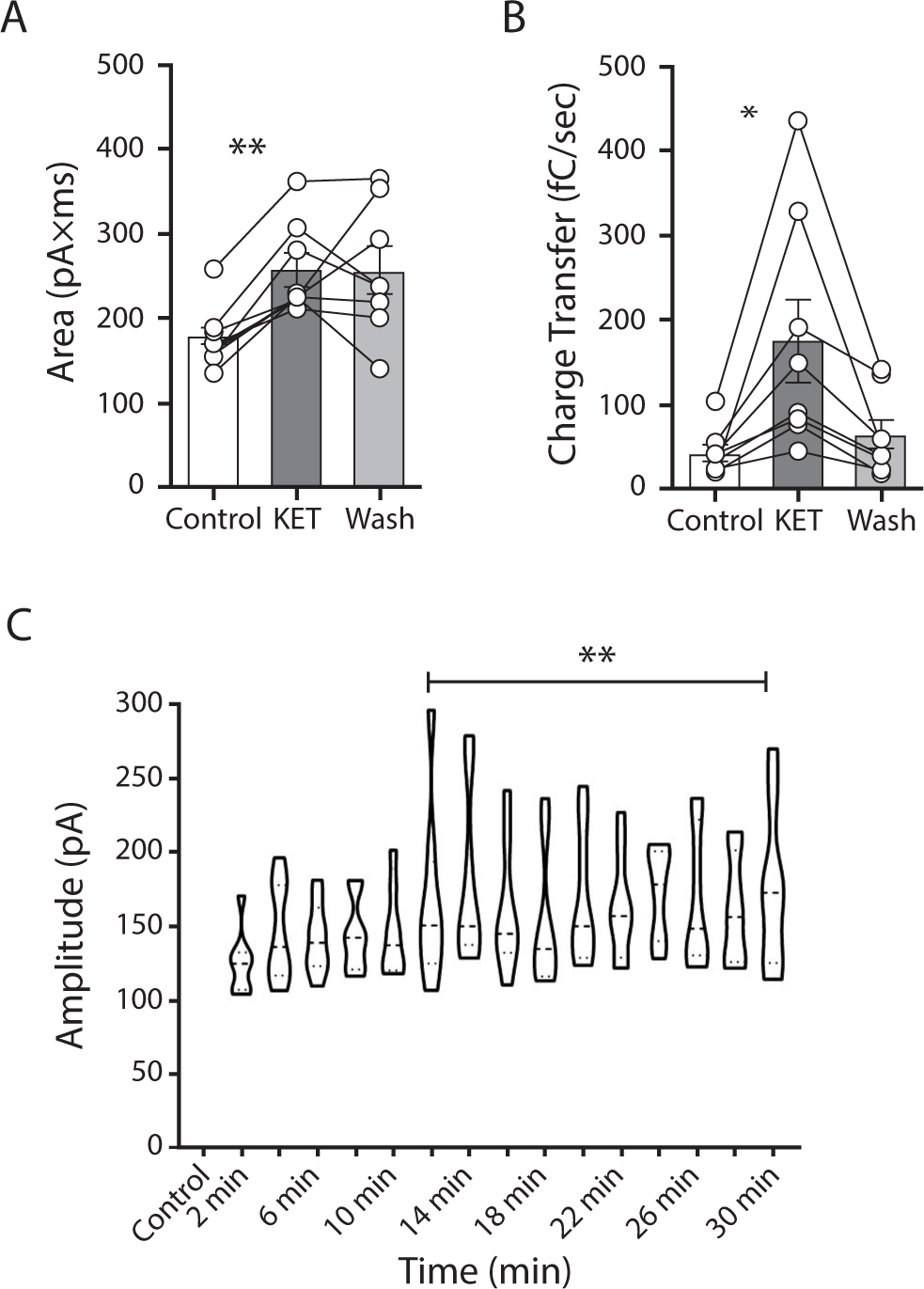
Effect of ketamine on the charge transfer and time course of the effect. A, B, Mean amplitudes of the average AUC and charge transfer (Q) of the events in control, during ketamine application and after wash-out. A, The AUC was significantly increased by the drug (*p* = 0.0048, n = 8, Friedman test). Dunn post hoc tests showed that the amplitude was significantly increased by ketamine (*p* = 0.0081). This effect was not reversible during wash-out (*p*>0.99). B, The charge transfer was also significantly increased by ketamine (one-way ANOVA: F (1.077, 7.541) = 8.818, *p* = 0.018, n = 8). Tuckey post hoc tests showed only a significant difference between the control and ketamine periods (*p* = 0.04). C, Violin plot of EPSC amplitude over time. A Kruskal Wallis test showed a significant increase of the amplitude over time (*p* = 0.0014, n = 8).

The average area under the curve (AUC) and the charge transfer (Q) of the events was determined and compared before, during ketamine application and after the washout (Fig. 3A, B). Both the AUC and Q were significantly increased by the drug (see details in the legend of Fig. 3A, B).

In other 5-HT neurons (n = 8), ketamine did not significantly affect sEPSC parameters (Fig. 4). Their amplitude was similar in control conditions and in ketamine (28.9 ± 0.8 vs 28.5 ± 1.2 pA, t = 0.57 with df = 7, *p* = 0.58, Student’s t test), as was their frequency (2.25 ± 0.47 vs 2.24 ± 0.47 Hz, t = 0.2263 with df = 7, *p* = 0.8274, Student’s t test). Corresponding plots are displayed in Extended Data Fig. 1-1. Rise times and decay times of the events were unaffected as well. These sEPSCs were also completely blocked by 10 µM CNQX.

**Figure 4.**
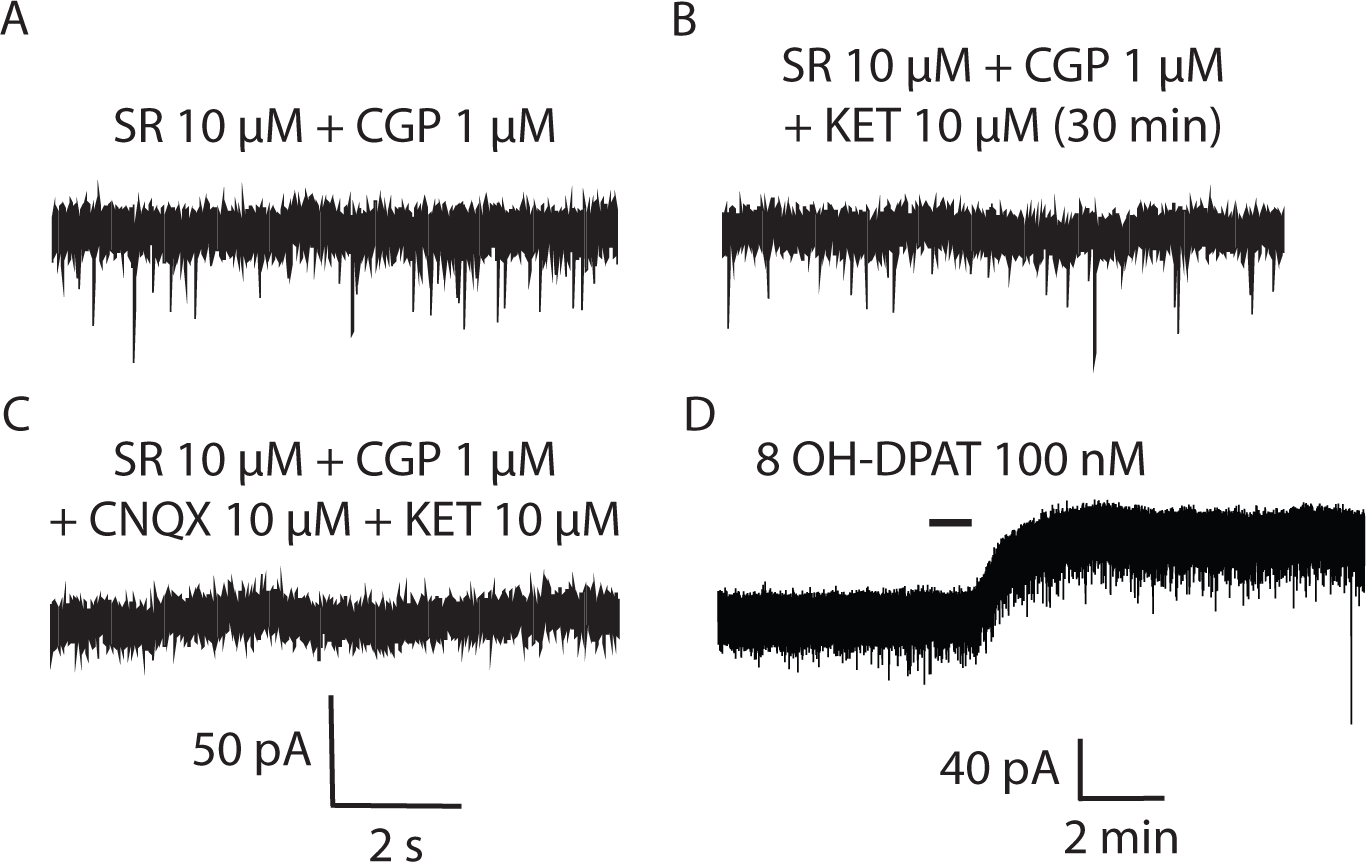
Another subgroup of pharmacologically identified 5-HT neurons is unresponsive to ketamine. Representative traces of sEPSCs recorded in the absence (A) and presence of ketamine 10 µM (B) in a (5-HT) neuron at a holding potential of −60 mV. C, the sEPSCs are blocked by CNQX (10 µM). D, The outward current elicited by the 5HT_1A_ agonist 8-OH-DPAT (100 nM) demonstrates the serotoninergic nature of the recorded neuron.

No criteria allowed ketamine-responsive and unresponsive neurons to be differentiated. Thus, the baseline frequency of the sEPSCs was similar in ketamine-responding and non-responding neurons (median values were 2.10 and 1.96 Hz in the responsive and non-responsive neurons respectively: Mann Whitney U test = 27, *p* = 0. 64, n = 16). Action potential waveforms and input resistances were also similar in both groups (input resistances: 382.5 ± 25.14 vs 411.2 ± 46.42 MΩ, t = 0.54 with df = 14, *p* = 0.59, unpaired t test, n = 16). In addition, the amplitude of the 8-OH-DPAT outward current was similar in both groups (82.2 ± 10.7 vs 71.8 ±7.3 pA (t = 0.8 with df = 14, *p* = 0.547, unpaired t test, n = 16).

We also recorded AMPA sEPSCs from non-5-HT neurons (n = 5). The frequency of these events was relatively low in these neurons in control conditions (Fig. 5). It was significantly lower than the frequency of sEPSCS in ketamine-responsive 5-HT neurons (median values were 1.10 and 2.10 Hz respectively (Mann Whitney U test = 4, *p* = 0.018, n = 13).

**Figure 5.**
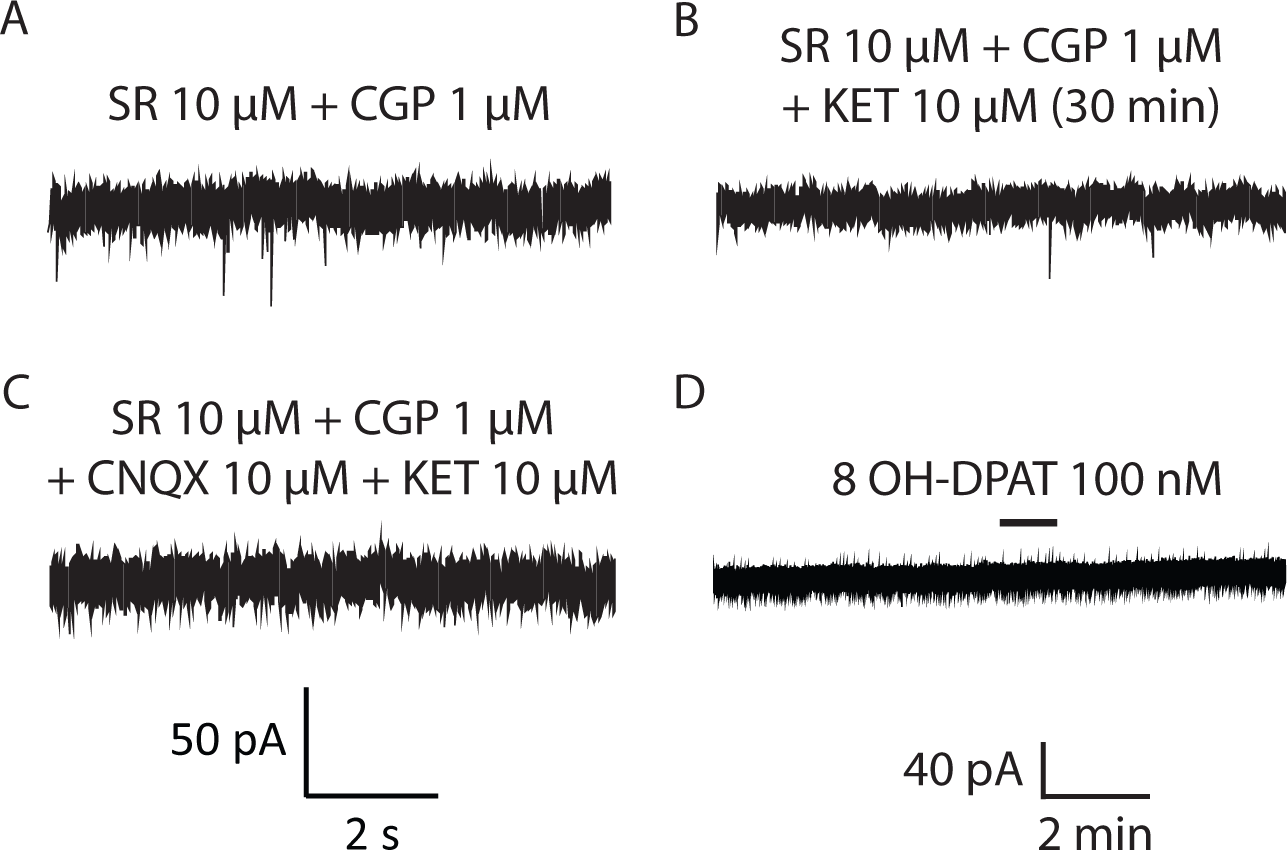
sEPSCs in a non 5-HT neuron of the DRN. A, B, Representative traces of spontaneous sEPSCs recorded in the absence (A) and presence of ketamine 10 µM (B). C, the sEPSCs are blocked by CNQX (10 µM). D, The 5HT_1A_ agonist 8-OH-DPAT (100 nM) has no effect in this neuron, demonstrating its non-serotoninergic nature.

Superfusion of ketamine did not affect any parameter of the sEPSCs in these neurons. The amplitude and frequency of the averaged sEPSCs in control and in the presence of ketamine were not significantly different. Rise times and decay times were similar as well. These sEPSCs were also completely blocked by 10 µM CNQX (Fig. 5C).

### Hydroxynorketamine mimicks the effect of ketamine, but has an overall weaker effect

The effects of 10 µM HNK were examined next. As shown in Figs. 6 and 7, the amplitude of the AMPA sEPSCs was significantly increased in about half of the recorded neurons by the drug. Out of 9 neurons, 6 neurons were responsive to HNK. This proportion is similar to the one found for its parent drug (z = 0.81, *p* = 0.42, two-sided Chi^2^ test). The amplitude of the events was increased in this population of cells (Fig. 7G; from 30.3 ± 2.3 to 41.2 ± 1.8 pA, t = 4.165 with df = 5, *p* = 0.0088, Student’s t test, n = 6). In the other 3 neurons the amplitude in the two conditions was similar (Fig. 8; median 26.26 and 25.81 pA, Wilcoxon test, *p* = 0.75, n = 3). In the responding neurons, the frequency of the events was also increased by the drug (from 1.74 ± 0.50 to 3.02 ± 0.69 Hz, t = 3.017 with 5 df, *p* = 0.029, Student’s t test, n = 6). It was unaffected in non-responding ones (Fig. 8; median: 1. 75 and 1.77 Hz, respectively, Wilcoxon test: *p* > 0.99, n = 3. Fig. 7 shows both the individual (A, B) and the average (C) cumulative probability plots for the responding neurons. A clear displacement to the right was observed for the amplitudes. The “EC_50_” value rose from 27.1 ±1.6 to 39.6 ± 2.8 pA (Fig.7D). Student’s t test showed that there was a significant difference between these values in the 2 groups (*p* = 0.017, n = 6). In addition, the cumulative probability distribution of the interevent intervals (Fig. 7D,E,F) was left shifted in ketamine in comparison to the control and the washout periods.The “EC_50_” value decreased from 500 to 100 ms. A Wilcoxon test did not show a significant difference, only trend, between the control and HNK periods (*p* = 0.0625, n = 8). In contrast the average interevent intervals were significantly decreased by the drug from 760 ± 142 to 475 ± 128 ms (not shown; Student’s paired t test, t = 4.17 with df = 5, *p* = 0.008, n = 6). In the 3 non-responsive neurons (Fig. 8), the interevent intervals varied from 576.1 to 563 ms (Fig. 8; *p* = 0.5, Wilcoxon test). Kinetic parameters of sEPSCs were unaffected in both responsive and non-responsive neurons.

**Figure 6.**
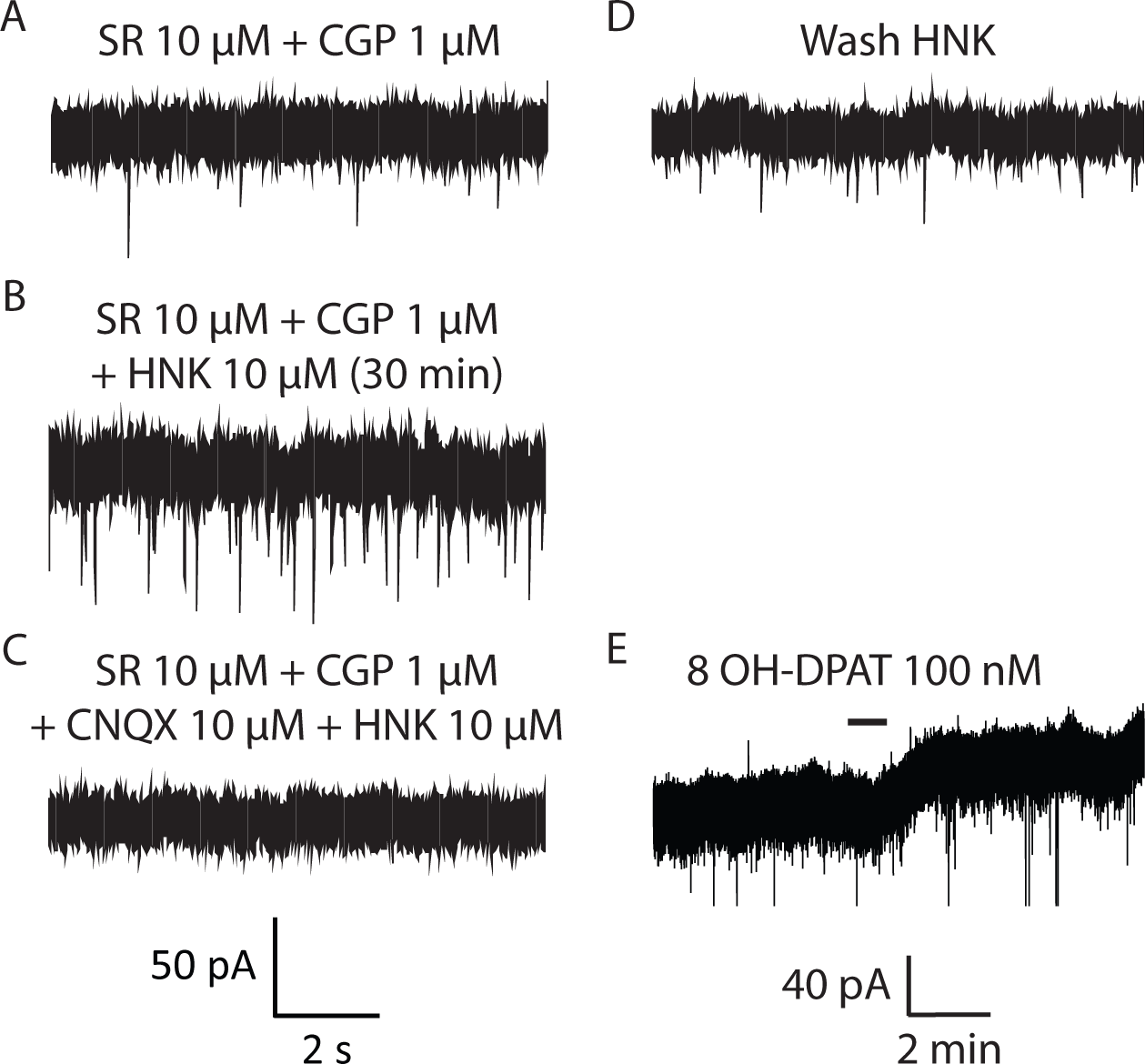
Effect of HNK on sEPSCs in a pharmacologically identified 5-HT neuron. A, B, Representative traces of sEPSCs recorded in the absence (A) and presence of HNK 10 µM (B). C, the sEPCs are blocked by CNQX (10 µM). D, sEPSCs recorded after the washout. (E) Effect of 8-OH-DPAT (100 nM).

**Figure 7.**
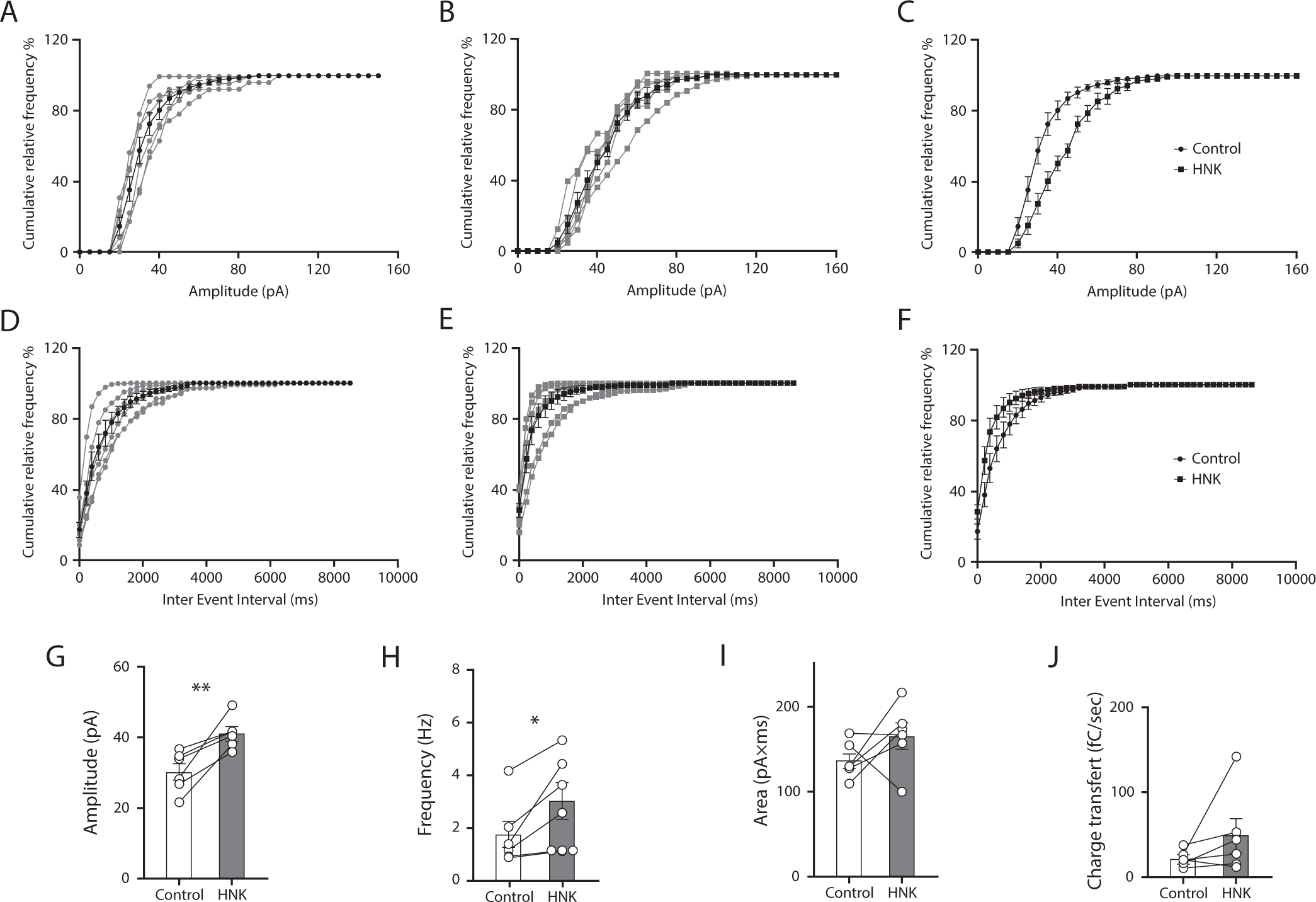
Quantification of the effect of HNK. A, B, Cumulative probability plots of sEPSC amplitude of the individual cells in control (A), and during ketamine (B) (Bin width = 5 pA, n = 8). C, Histogram of the averaged data for the amplitude. C, The cumulative probability plots from all the cells were averaged to generate the cumulative probability plots of sEPSC amplitude in each condition (control, and HNK). A clear displacement to the right was observed for the amplitudes. D, E, Cumulative probability of sEPSCs interevent interval of the individual cells in control (D) and during HNK (E) (Bin width = 200 ms). F, Averaged cumulative probability plots. G, H, The histograms represent the averaged data for the amplitude and frequency. I, J, AUCs and charge transfer (Q) of the events before, during and after ketamine application. I, The AUC was not significantly increased by the drug (t = 1.386 with df = 5 *p* = 0.22, n = 6, Student’s paired t test). G, The charge transfer was not significantly increased either by ketamine (t = 1.15, df = 5 *p* = 0.19, n = 6, Student’s paired t test).

**Figure 8.**
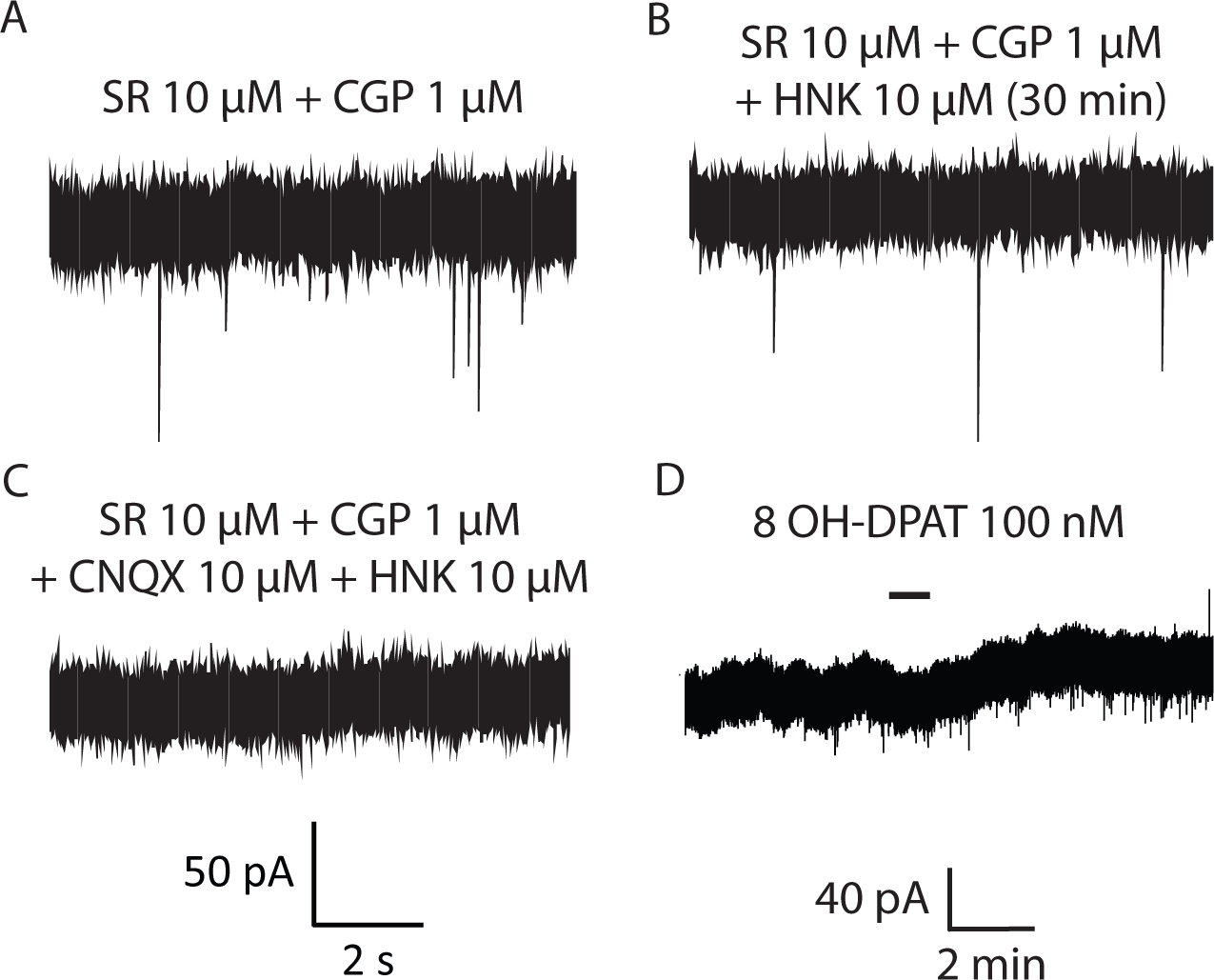
HNK has no effect on a subgroup of pharmacologically identified 5-HT neurons. A, B, Representative traces of sEPSCs recorded in the absence (A) and presence of HNK 10 µM (B). C, the sEPSCs are blocked by CNQX (10 µM). D, The outward current elicited by the 5HT_1A_ agonist 8-OH-DPAT (100 nM) demonstrates the serotoninergic nature of the recorded neuron.

### Mechanism of the ketamine potentiation of AMPA sEPSCs

In order to gain insight into the mechanism of sEPSC potentiation by ketamine in 5-HT neurons, we first asked whether its effect involved its canonical action as a NMDA antagonist. For this purpose, we studied its effect in the continuous presence of 50 µM D-AP-5 (n = 7). An example is shown in Figure 9.In these conditions, ketamine had no effect on either the amplitude or the frequency of the events. The results are summarized in Fig. 11A and B. Of note, the control sEPSC amplitude tended to be higher in the continuous presence of APV than in its absence (Fig. 9E; 33.8 ± 2.0 vs 28.7 ± 1.5 pA, t = 2.04 with df = 13, *p* = 0.06, Unpaired t test, n = 15), strongly suggesting the possibility of an at least partial occlusion phenomenon. Kinetic parameters of the sEPSCs were unaffected as well by ketamine in these conditions. The relevant cumulative probability plots are shown in Extended Data Fig. 1-2. Thus, the antagonistic action of ketamine on NMDA channels seems important for the AMPA sEPSC potentiation that it induces.

**Figure 9.**
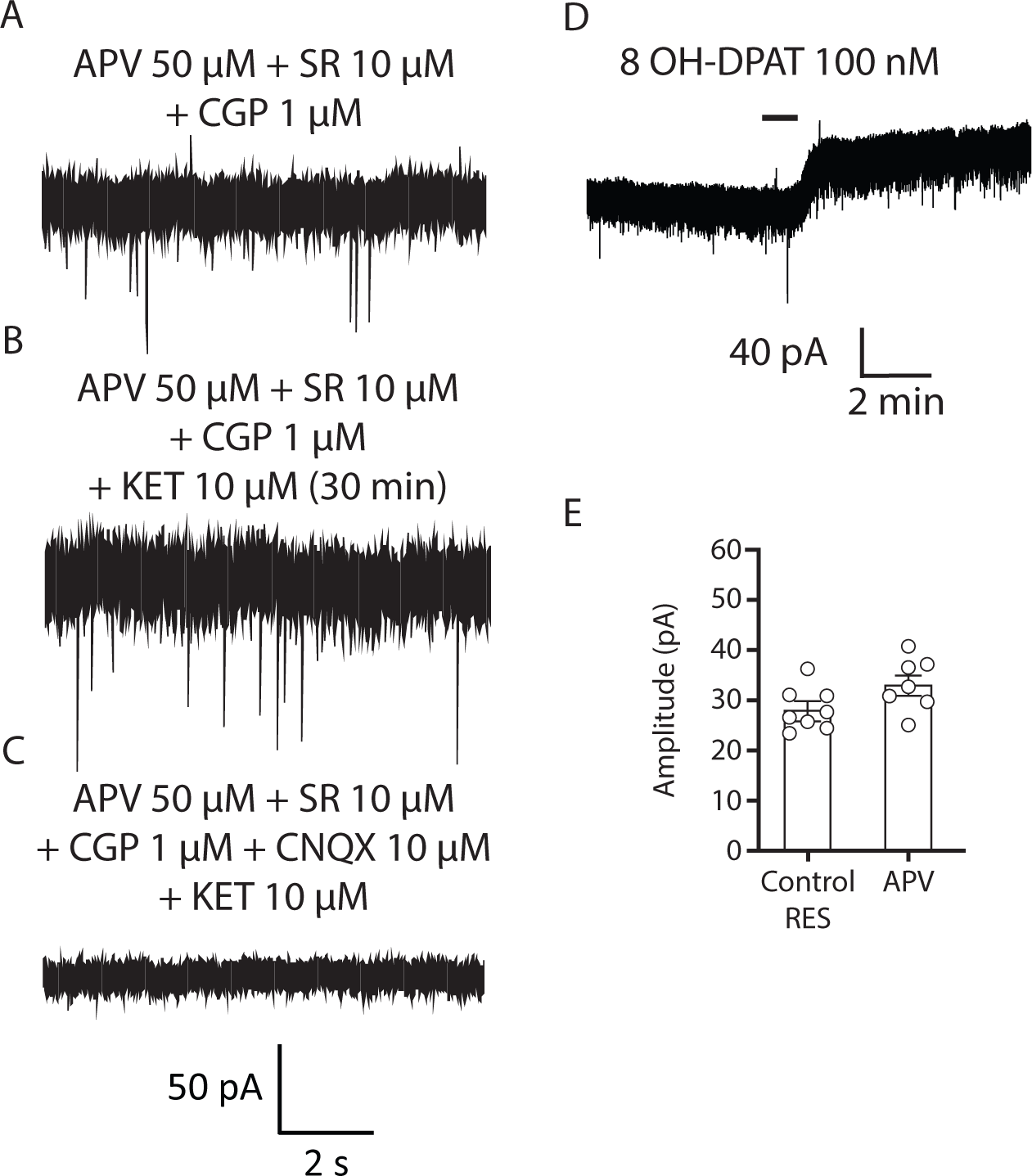
Ketamine is devoid of any effect on the amplitude and frequency of sEPSCs in the presence of 50 µM D-AP-5. A, B, Representative traces of sEPSCs recorded in the absence (A) and presence of HNK 10 µM (B). C, the sEPSCs are blocked by CNQX (10 µM). D, Outward current elicited by the 8-OH-DPAT (100 nM) in the same neuron. E, The control sEPSC amplitude tended to be higher in the continuous presence of APV than in its absence.

We next wondered whether ketamine’s effect would persist in the continuous presence of 1 µM TTX (Fig. 10), which abolishes action potential dependent neurotransmitter release. In these conditions, the mean frequency and amplitude of the control miniature EPSCs (mEPSCs) (Fig. 11C, and D) were 2.83 ± 0.86 Hz and 29.90 ± 3.01 pA, respectively (n = 7). These parameters were actually not different from those observed in control conditions (i.e. in the absence of the toxin). We did not see any difference after superfusion with ketamine, neither in terms of frequency (actually a trend towards a *decrease* in frequency was observed: from 2.83 ± 0.86 to 2.48 ± 0.74 Hz; t=2.28 with df = 6, *p* = 0.06, Student’s t test) or in terms of amplitude (29.9 ± 3.0 to 29.0 ± 2.7 pA; t = 2.20 with 6 df, *p* = 0.069, Student’s t test) (Fig. 11C, D).

**Figure 10.**
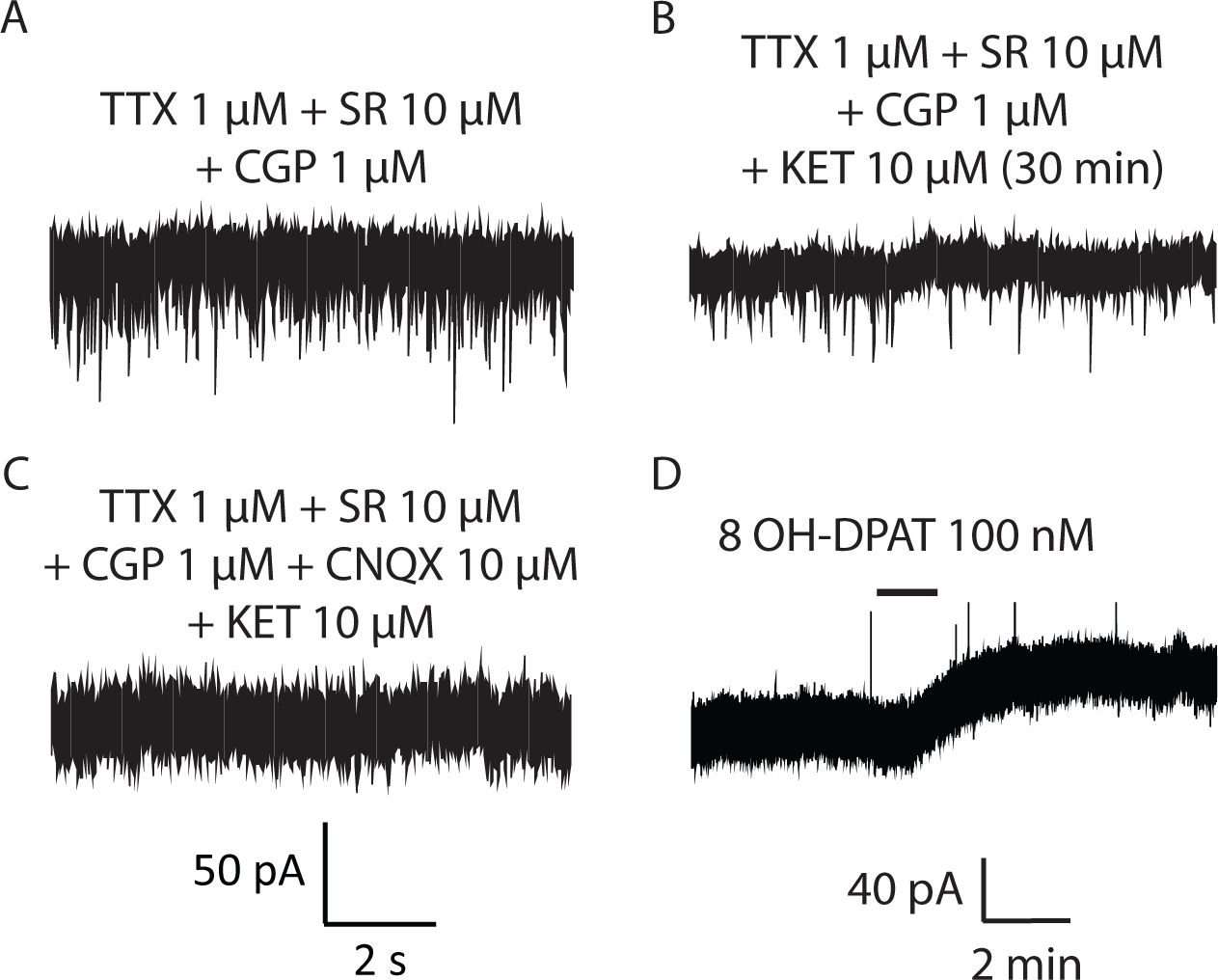
The effect of ketamine was not observed in the presence of tetrodotoxin (TTX). A, B, Representative traces of mEPSCs recorded in the absence (A) and presence (B) of ketamine 10 µM. C, The mEPCs are blocked by CNQX (10 µM). (D) Outward current elicited by the 8-OH-DPAT (100 nM).

**Figure 11.**
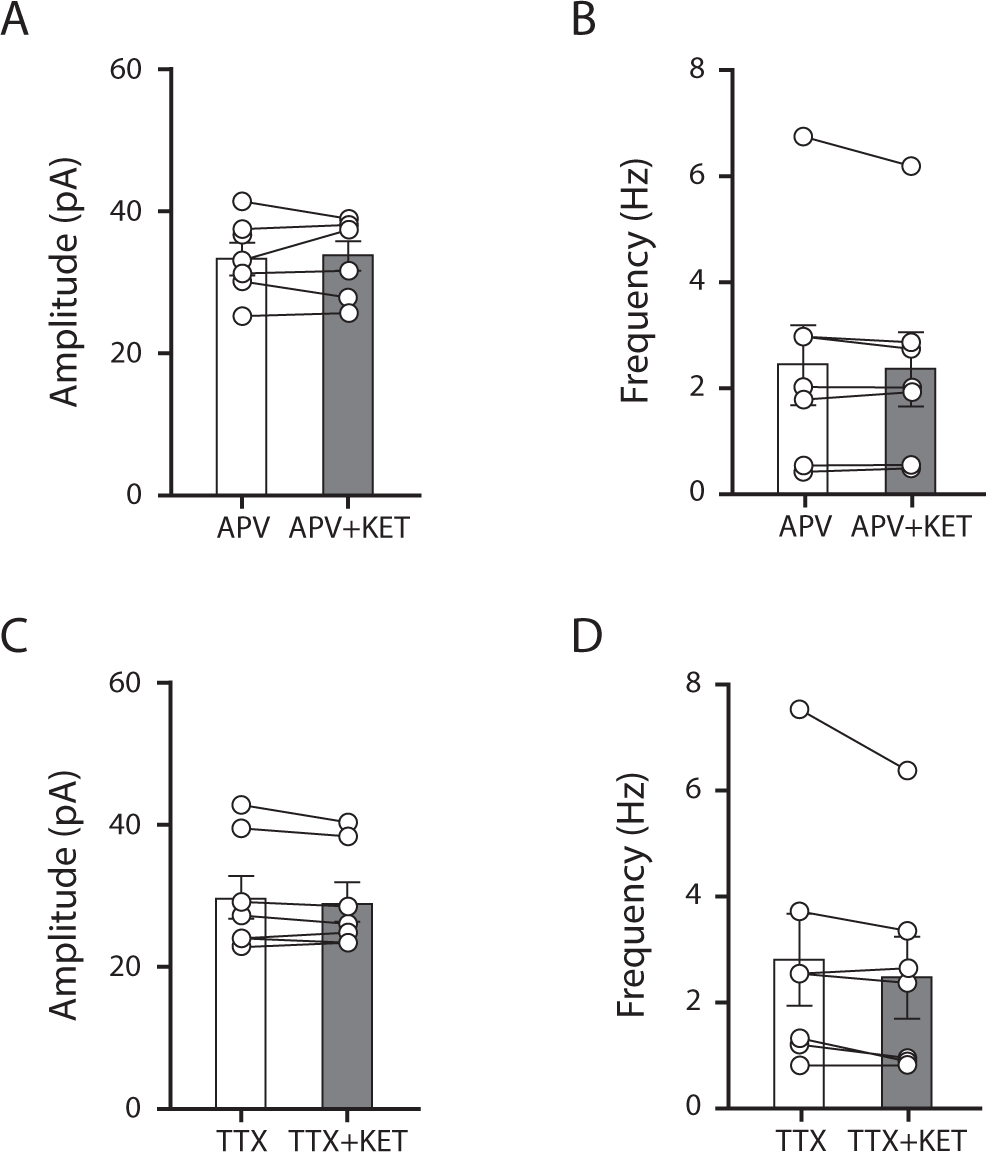
Histograms of the average amplitude and frequency of the events in the experiments performed in the presence of D-AP-5 or TTX. A, amplitude of the events under D-APV did not change during ketamine superfusion (33.3 vs 37.7 pA, *p* = 0.68, Wilcoxon test, n = 7). B, Frequency of sEPSCs was unaffected by ketamine 10 µM (2.4 ± 0.8 vs 2.3 ± 0.7 Hz, t = 1.019 with df = 6, *p* = 0.34, Student’s t test). C, In TTX, the mean amplitude of the mEPSCs was not modified by ketamine 10 µM (29.9 ± 3.0 pA to 29.1 ± 2.71 pA, t = 2.20 with df = 6, p = 0.069, Student’s t test, n = 7). D, Lack of effect of ketamine on the frequency of the events in TTX (2.8 ± 0.9 vs 2.5 ± 0.7 Hz, t = 2.28 with df = 6, *p* = 0.062, Student’s t test, n = 7).

On the other hand, the decay time increased in the presence of ketamine from a median value of 1.50 to 1.70 ms (Wilcoxon test: *p* = 0.0313, n = 7), but the rise time was unaffected (median values were 0.49 and 0.57 ms, respectively Wilcoxon test: *p* = 0.15, n = 7). These results show that ketamine’s potentiation of AMPA EPSCs vanishes when action potential propagation is blocked. The relevant cumulative probability plots are shown in Extended Data Fig. 1-3. These mEPSCs were completely blocked by 10 µM CNQX as well.

## DISCUSSION

Ketamine has previously been shown to affect the excitability of neurons located in many brain regions, including prefrontal cortex, hippocampus and habenula (Li et al., 2010; Zanos et al., 2016; Yang et al., 2018). In general, an increase in AMPA receptor density is considered to be an important final common pathway (probably following BDNF release), although some authors consider that NMDA blockade is necessary and sufficient to explain ketamine’s fast antidepressant action (Yang et al., 2018; Widman and McMahon, 2018).

Results from different research groups also point to the possibility of 5-HT involvement in ketamine’s action (du Jardin et al., 2016), including *in vivo* evidence for an action of ketamine on the excitability of 5-HT neurons via activation of AMPA and nicotinic receptors (Nichitani et al., 2014). A very recent mouse brain slice study suggested that a high concentration of ketamine (50 µM) promotes activation of AMPA receptors in the dorsal raphe (Llamosas et al., 2019). However, the neurochemical nature of recorded cells was not clear from their results, limiting our ability to interpret the data.

Using a rigorous identification of the recorded neurons, we show here that both ketamine and it’s probably active metabolite HNK increase AMPA receptor activation in a subset (~50%) of 5-HT neurons. Both ketamine-sensitive and insensitive neurons are clearly serotonergic, since their 5-HT_1A_ tone is similar. To our knowledge, this is one of the first demonstrations of a heterogeneous effect of a compound within the population of DRN 5-HT neurons. For example, effects of the activation of alpha_1_ noradrenergic receptors, orexin receptors and 5-HT_1A_ receptors are observed in most 5-HT neurons in this region (Pan et al, 1994; Ishibashi et al 2016). Because inputs and outputs of these neurons are segregated, in particular in terms of outputs to the prefrontal cortex and amygdala (Ren et al., 2018), it is possible that ketamine is able to bias 5-HT transmission towards the PFC-projecting neurons, which might promote active coping and decrease anxiety. Experiments on 5-HT neurons retrogradely labelled after injection of fluorescently labeled latex beads in one or the other region will allow us to test this hypothesis.

One question that arises is how the effect of ketamine and HNK can be so dichotomous within the DRN. In this respect, we found that the effect of ketamine is much attenuated in the presence of TTX, suggesting that the local network is important in its action and that presynaptic effects are involved. It may be that the synaptic micro-environment of 5-HT neurons is heterogeneous in this area. For example, glutamate terminals innervating ketamine-responsive and unresponsive 5-HT neurons might originate from different areas and have differential properties, e.g. in terms of heteroreceptors and second messenger pathways. In addition to the main inputs to the region (prefrontal cortex, lateral habenula and hypothalamus), one input could be provided within the DRN itself, since a subpopulation of 5-HT neurons has been shown to express a vesicular glutamate transporter (VGLUT3) (Gras et al., 2002) and to release glutamate (Liu et al., 2014). This will be the subject of further investigation in our laboratory.

The time course of the effect of ketamine was clearly slower than the one of direct receptor agonist such as 8-OH DPAT (compare Figs 1E and 3C), suggesting that it induces a cascade of events that is rather complex. Our experiments did not address the nature of the second messenger pathway that could be implicated in the AMPA sEPSC potentiation by ketamine. Similar time courses for the effect of the drug have been found previously in slices (e.g. Zanos et al., 2016).

We found that superfusion of the competitive NMDA antagonist APV increased the mean amplitude of the sEPSCs from 28.7 to 33.8 pA. In the absence of APV, this amplitude rose from 28.7 to 46 pA in the ketamine-sensitive neurons. These results suggest that a partial occlusion of the effect of ketamine may have been produced by APV. However in the absence of APV, the mean increase of amplitude produced by ketamine in responsive neurons was quite larger. How can we interpret this? One possibility is that the non-competitive nature of ketamine’s action on NMDA receptors is somehow able to modulate the target receptors in a different way than APV.

The robust potentiation of AMPA transmission that we observed in a subpopulation of 5-HT neurons is likely to translate into increases in firing rates in vivo. This should be tested using telemetric recordings in awake animals. It will also be important in the future to test *in vivo* whether the drug does induce a bias towards activation of PFC-projecting 5-HT neurons by performing multi-unit Ca^2+^ transient recordings in neurons with a known projection, using a combination of GCaMP transfection and Cre-Lox technology.

In conclusion, ketamine, and to a lesser degree HNK, are able to potentiate AMPA transmission in a subset of pharmacologically identified DRN 5-HT neurons, which may contribute to their acute antidepressant effect.

## Acknowledgements

Funded by grant FSR-S-SS-18/23 of the « Fonds spéciaux de la recherche » from Liège University (2018). CH has received a « bourse d’excellence » from « Wallonie Bruxelles International » to support her salary for 4 years. We thank all members of the Neurophysiology laboratory, as well as Dr. G. Morelli, for helpful discussions. We are especially thankful to Laurent Massotte for his invaluable help with the artwork.

## Extended data

**Fig 1-1.**
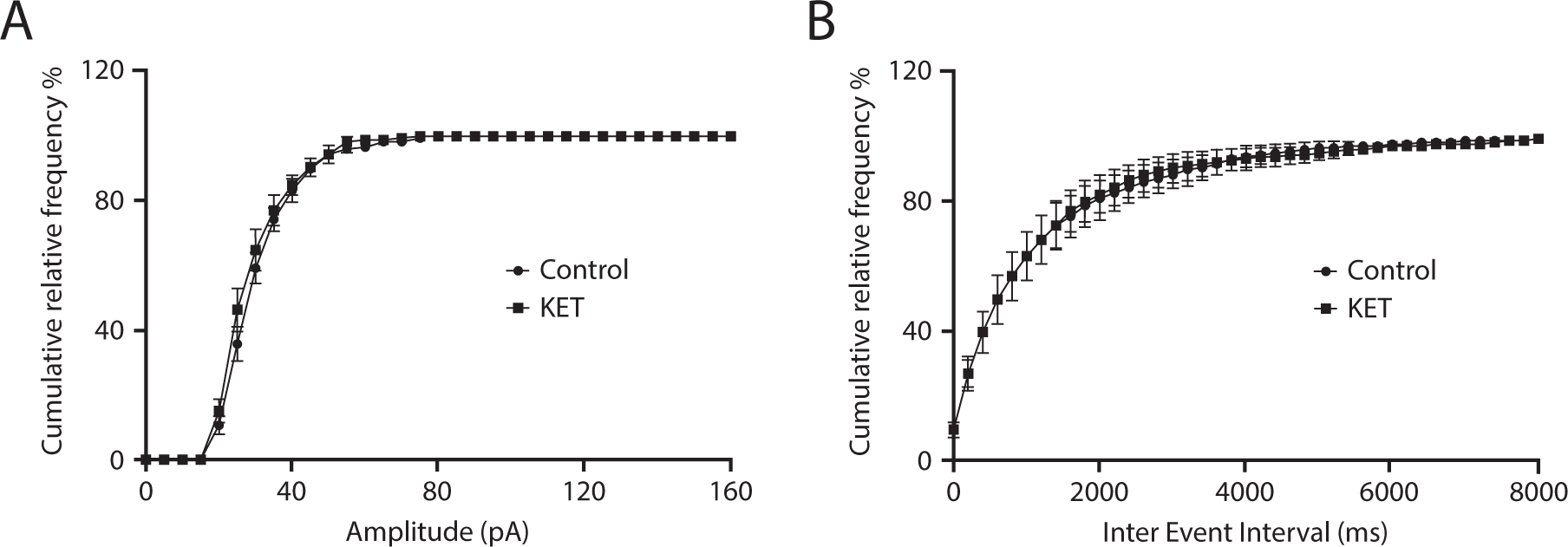
A subpopulation of pharmacologically identified 5-HT neurons are unresponsive to ketamine. A, B The cumulative probability plots from all the cells were averaged to generate the cumulative probability plots of sEPSC amplitudes (A) and interevent intervals (B). (Bin width = 5 pA). The “EC50” value of the amplitude was 27.8 ± 1.2 and 27.8 ± 1.4 pA in control and ketamine, respectively (t = 0.000 with df = 7, *p*>0.99, Student’s t test, n = 8). B, Averaged cumulative probability plots of interevent interval in control, and during ketamine 10 µM (Bin width = 200 ms, n = 8). “EC_50_” values were 800 ± 191 and 812 ± 198 ms (t = 0.35 with df = 7, *p*=0.73, Student’s t test, n = 8).

**Figure 1-2.**
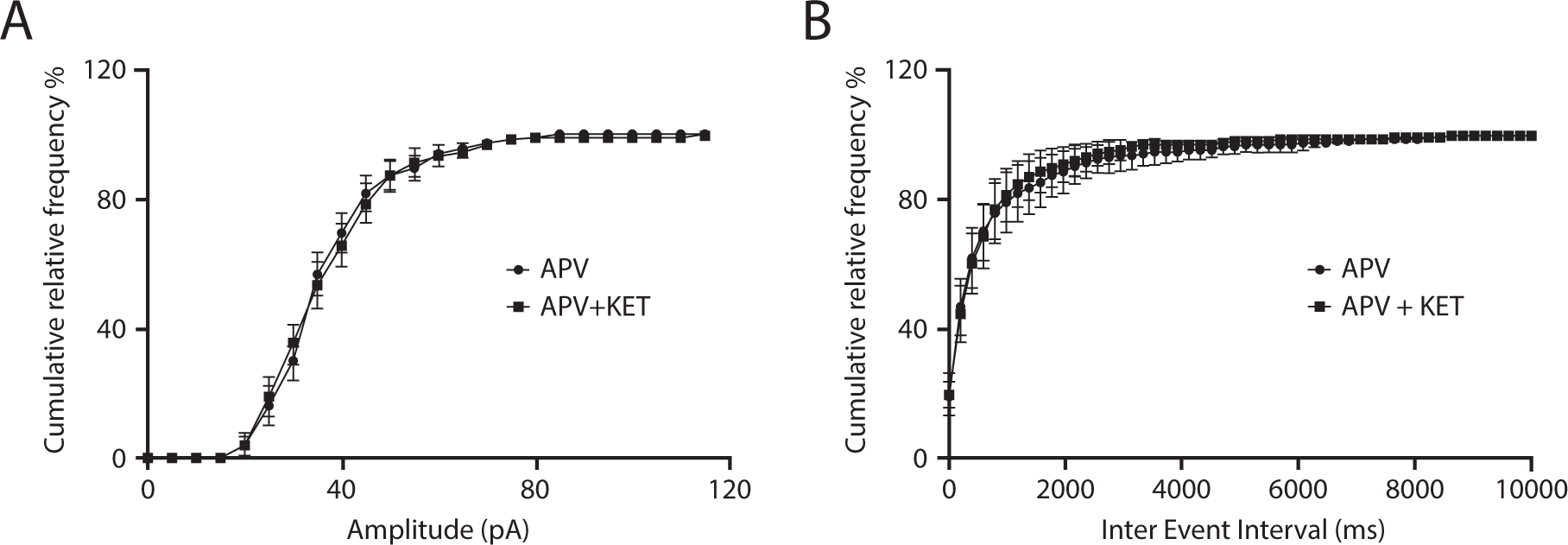
Ketamine was devoid of any effect on the amplitude and frequency of sEPSCs in the presence of 50 µM D-AP-5. A, Averaged cumulative probability plots of sEPSCs amplitude in the presence of D-APV-5 in control conditions and during ketamine 10 µM (Bin width = 5 pA). The “EC50” value of the amplitude was 34.6 ± 2.2 to 33.6 ± 2.2 pA, (t = 2.12 with df = 6, *p* = 0.0781, Student’s t test n = 7). B, Same for the intervent intervals (Bin width = 200 ms). The median value of the “EC50” of the interevent intervals was 300 ms in both conditions (*p* = 0.5, Wilcoxon test, n = 7).

**Figure 1-3.**
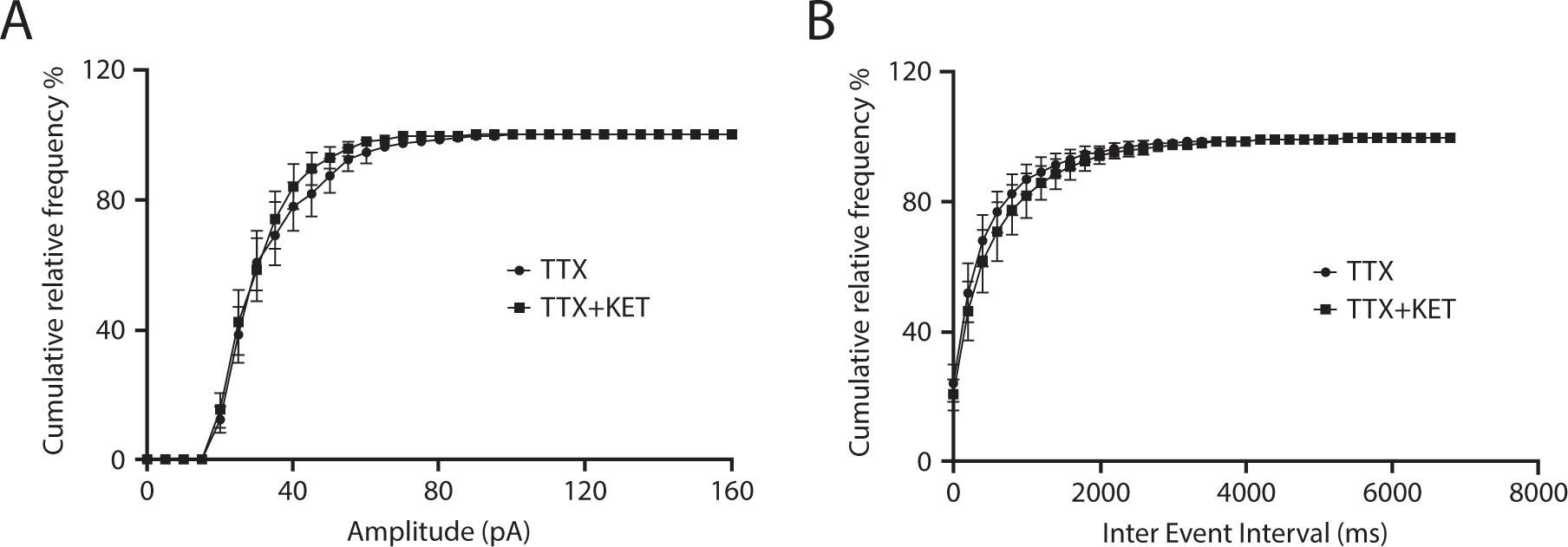
No effect of ketamine was observed in the presence of tetrodotoxin (TTX). A, Averaged cumulative probability plots of mEPSCs amplitude in control and during ketamine 10 µM (Bin width = 5 pA). The “EC50” value of the amplitude was 29.6 ± 3.6 and 28.6 ± 2.4 pA, (t = 0.62 with df = 6, *p* = 0.5, Student’s t test n = 7). B, Same for the intervent intervals (Bin width = 200 ms). The median value of the “EC50” of the interevent interval was 100 ms in both conditions (*p* = 0.2, Wilcoxon test, n = 7).

